# Mimivirus encodes an essential MC1-like non-histone architectural protein involved in DNA condensation

**DOI:** 10.1101/2024.02.22.580433

**Authors:** Deepti Sharma, Fasséli Coulibaly, Kiran Kondabagil

## Abstract

The first giant virus discovered, *Acanthamoeba polyphaga* mimivirus (APMV), has a 1.2 Mb dsDNA genome organized as genomic fiber within the capsid. This fiber is comprised of a proteinaceous shell of ∼30 nm diameter that encloses the folded DNA. Surprisingly, for the assembly of the enormous genome of APMV, no DNA condensing protein has been reported to date. Our analysis of the uncharacterized packaged protein complement of Mimivirus led to the identification of a putative DNA-bending archaeal MC1 domain in a hypothetical protein (gp275) coded by the R252 gene. Gene knock-out analysis shows that gp275 is critical for viral multiplication. Biochemical and microscopic characterization further demonstrates the compaction of DNA upon binding to gp275. Together, this study suggests that gp275 is an MC1-like architectural protein involved in the organization of the genomic DNA within the capsid of Mimivirus.

## Main

Genome organization is a fundamental phenomenon in the three domains of life aided by DNA-binding proteins. The physiological state of genomic DNA as an organized entity is associated with genome function and gene regulation. In eukaryotes, the genome is arranged into chromatin structure with histone proteins associated with DNA^1,2,3^. The bacterial genome is organized into condensed conformations by nucleoid-associated proteins (NAPs) such as H-NS, IHF, Fis, and HU^4,5^. Chromatin proteins in archaea mainly include histones and other architectural proteins such as Alba or MC1, Methanogen Chromosomal protein 1^6,7^. These DNA-binding proteins help in condensing the genome by wrapping, bending or bridging DNA and further assisting cellular processes such as DNA replication and gene transcription^8^.

The discovery of the association of viral genome with histone-like proteins in the past two decades has advanced our understanding of genome organization in viruses. Some viruses encode a single histone like bracoviruses encoding the H4 gene^9^ and pandoraviruses encoding H2B^10^. The members of some families belonging to the group, Nucleocytoplasmic Large DNA Viruses (NCLDVs), encode homologs of eukaryotic histones. The NCLDVs group is comprised of dsDNA viruses with larger genomes and capsids compared to most other viruses and have been classified into phylum, *Nucleocytoviricota*^11,12,13,14^. An allometric relationship between capsid volume and genome size has been observed in the case of NCLDVs. The enormous genome length of these viruses is associated with a large number of proteins encoded by the viral genome^15,16^. To accommodate both large genomes and a considerable number of packaged proteins, the organization of genomic DNA in a compacted form inside the capsid may occur as an expected phenomenon.

The *Marseilleviridae* family encode four histone proteins as doublets of H2A-H2B and H3-H4^17^. These histone doublets have been shown to form nucleosomes upon *in vitro* assembly^18^. While Iridoviruses code for H3-H4 histone doublet^19^, Medusavirus encode all five eukaryotic histones^20^. Thus, the pervasive presence of histone-like genes in various families of *Nucleocytoviricota* implies their potential role in the viral genome packaging.

*Acanthamoeba polyphaga* mimivirus, the prototype member of the *Mimiviridae* family of *Nucleocytoviricota*, has an enormous dsDNA genome of 1.2 Mb and a giant capsid size of ∼500 nm^21,22^. The Mimivirus genome encodes nearly a thousand proteins^23^. About half of the predicted genes are ORFans^24,25^ including the majority of the 200 proteins incorporated in mature viral particles^31,32^. The Mimivirus capsid contains an inner nucleoid or nucleocapsid formed by a membrane-delimited compartment of ∼340 nm diameter encasing the viral genome and several proteins ^26,27,28,29,30^.

Atomic force microscopy study of the Mimivirus infection cycle showed the organization of the viral genome in a condensed form along with proteins^33^. Recent cryo-electron microscopy (Cryo-EM) and cryo-electron tomography (Cryo-ET) studies of opened capsids have revealed the release of the Mimivirus DNA as genomic fibers. The fiber accommodates the genome in a proteinaceous shell, which is composed predominantly of two GMC (glucose-methanol-choline)-oxidoreductases that lead to the speculation of a role for them in packaging the genome^34^. However, the laboratory-generated Mimivirus mutant-M4 showed the absence of both GMC-oxidoreductases in the proteomics data. These mutated Mimivirus particles could package DNA inside the capsid and retain infectivity similar to wild-type Mimivirus^35^. Furthermore, the gene knockout analysis of both GMC-oxidoreductases has further established that they are not essential for genomic fiber formation and that no fitness cost is associated with their deletion^36^. Despite being one of the first giant viruses discovered with a genome of substantial size, how the Mimivirus genome is compacted and packaged inside the virion is poorly understood.

In this study, we report the identification and characterization of an MC1-like DNA condensing protein in Mimivirus. Sequence analysis of all virus-coded proteins incorporated into the virion helped us identify a putative MC1 domain in the Mimivirus gp275 (encoded by gene R252), previously annotated as a hypothetical protein. MC1 (Methanogen Chromosomal protein 1) is a DNA-bending protein in methanogenic archaea, which is involved in genome organization^37, 38, 39^. The sequence analysis and phylogeny of gp275 with archaeal MC1 protein suggests gp275 to be an architectural protein of Mimivirus that is involved in compacting DNA and thus may help in genome packaging. We show that gp275 is an oligomeric protein that binds and bends the DNA. Furthermore, R252 gene knockout experiments suggested that it is an essential gene with its involvement in the mimiviral replication cycle. We also demonstrate the presence of gp275 in the virion by mass spectrometry and microscopy. In sum, we show that gp275 is an essential non-histone protein that might be involved in the chromatinization of Mimivirus genomic DNA.

## Results

### Mimivirus gp275 is distantly related to archaeal non-histone chromosomal protein

To identify the potential DNA architectural protein/s in Mimivirus, we analyzed the mimiviral hypothetical protein sequences available in the NCBI database. The domain search using InterProScan led to the identification of a putative MC1 domain in the *Acanthamoeba polyphaga* mimivirus gp275 protein. MC1 is a monomeric non-histone architectural protein present in the methanogenic archaea, *Euryarchaea,* which protects DNA against thermal denaturation^37,40,41^ and helps in organizing DNA by inducing bends^38,39^. The PSI blast search against the *Mimiviridae* family revealed the presence of gp275 homologs in all three lineages (A, B and C) of the viral family, including the unclassified mimiviruses. The homologs of Mimivirus gp275 were observed in two other families of the phylum *Nucleocytoviricota*, including members of the family *Phycodnaviridae* (*Paramecium bursaria* Chlorella virus 1 and *Acanthocystis turfacea* Chlorella virus) and metagenome-assembled Marseillevirus. Mimivirus gp275 showed ∼ 30% identity with MC1 protein sequences of archaea (Fig. 1a). Modelling of the 3 dimensional structure with AlphaFold2^44,74^ revealed the conservation of the global fold of the MC1 protein of archaea in Mimivirus gp275 (Fig. 1b) as well as other viruses shown in Fig. 1a. The fold is a pseudo barrel which consists of five anti-parallel beta strands and one alpha helix (β1-β2-α1-β3-β4-β5) all connected by loops^42^. The gp275 homologs among all three mimiviral lineages and tupanviruses showed the presence of a lysine residue within the α-helix in place of the arginine residue of the archaeal MC1 protein critical for DNA bending^43^. The other unclassified mimiviruses, such as *Aureococcus anophagefferens* virus and *Cafeteria roenbergensis* virus BV PW1 from group II, along with Marseillevirus and phycodnaviruses, showed the presence of serine, tyrosine, methionine, or glycine, respectively, in the place of the critical arginine/lysine residue at that position.

**Fig. 1.**
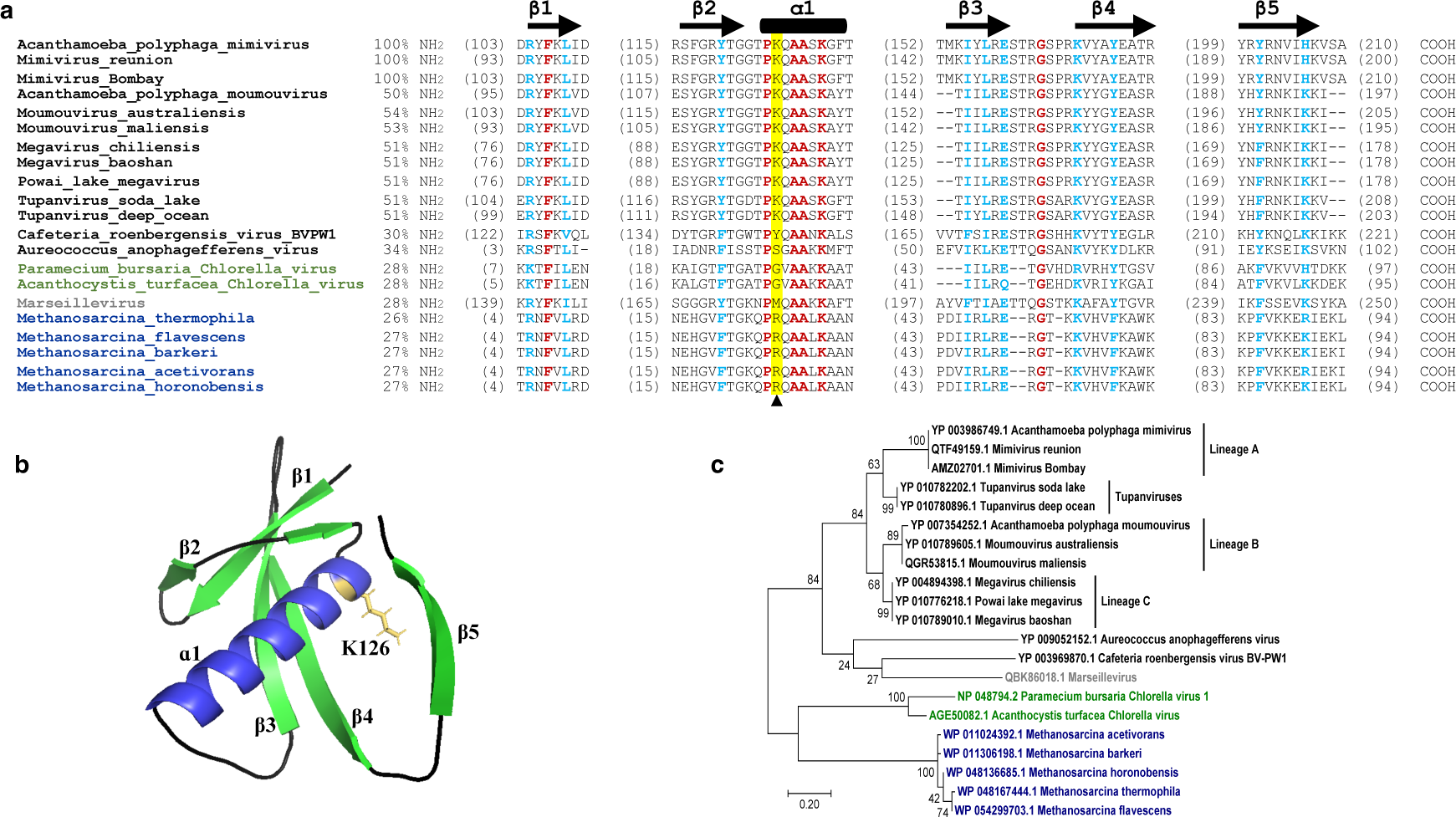
Amino acid sequence analysis of Mimivirus gp275 showing the presence of MC1 domain. **a,** gp275 multiple sequence alignment was created using ClustalW^46^. Sequences used from archaea belong to the genus *Methanosarcina* (WP_048167444.1, WP_054299703.1, WP_011306198.1, WP_011024392.1 and WP_048136685.1) and; from mimiviruses: *Acanthamoeba polyphaga* mimivirus (YP_003986749.1), Mimivirus reunion (QTF49159.1) and Mimivirus Bombay (AMZ02701.1) of lineage A; *Acanthamoeba polyphaga* moumouvirus (YP_007354252.1), Moumouvirus australiensis (YP_101789605.1) and Moumouvirus maliensis (QGR53815.1) of lineage B; Megavirus chiliensis (YP_004894398.1), Megavirus baoshan (YP_010789010.1) and Powai lake megavirus (YP_010776218.1) of lineage C; Tupanvirus soda lake (YP_010782202.1) and Tupanvirus deep ocean (YP_101780896.1) from tupanviruses; *Cafeteria roenbergensis* virus BVPW1 (YP_003969870.1) and *Aureococcus anophagefferens* virus (YP_009052152.1) from unclassified mimiviruses. *Paramecium bursaria* Chlorella virus 1 (NP_048794.2) and *Acanthocystis turfacea* Chlorella virus (AGE50082.1) from phycodnaviruses; and Marseillevirus (QBK86018.1) sequence from metagenome-assembled genome (Marseillevirus LCMAC101). *Mimiviridae* family members are highlighted in black, *Phycodnaviridae* in green, Marseillevirus in grey and archaea in dark blue. The secondary structure conservation with the MC1 domain is shown with five beta sheets (β1-β5) and one alpha helix (ɑ1). Color coding: red-identical residues; blue-highly similar residues in a group. Numbers before aligned sequences indicate the percentage similarity and numbers in parenthesis indicate the position of amino acid in the sequence. The arrowhead indicates the position of arginine residue critical for DNA-bending in archaea and conserved lysine in mimivirus lineages (highlighted row in yellow). **b,** gp275 MC1 domain monomer structure predicted by AlphaFold2 excluding disordered N and C-terminal regions. Color is shown as green-beta sheets (β1-β5), blue-alpha helix (ɑ1), black-turns/loops and yellow-conserved Mimivirus lysine (K126) residue in ɑ1. **c,** Maximum likelihood phylogenetic tree of Mimivirus gp275 homologs from NCLDVs with MC1 protein sequences of archaea. *Mimiviridae* family members are highlighted in black; *Phycodnaviridae* in green; Marseillevirus in grey and archaea in blue. Lineages A, B and C of *Mimiviridae* along with tupanviruses are labeled. The scale bar represents the number of substitutions per site.

The phylogenetic analysis further revealed the association of Mimivirus gp275 with other NCLDVs homologs and archaeal MC1 protein (Fig. 1c). While the gp275 homologs from *Mimiviridae* lineages A, B and C along with tupanviruses clustered together, the other unclassified mimiviruses formed a separate clade with metagenome-assembled Marseillevirus. Furthermore, the gp275 homologs of the *Phycodnaviridae* family were observed to be more closely related to archaeal proteins than mimiviral MC1 proteins. Based on the phylogenetic analysis, the most parsimonious explanation of MC1 gene acquisition by mimiviruses could be the lateral transfer of genes from archaea to an algal ancestor (no algal hits of MC1 protein were found during the BLASTp search) inhabiting the same niche through symbiotic association^47^ and then to viruses having algal hosts followed by their diversification to infecting protozoans^48,49,78^. The phylogenetic and sequence analysis of gp275 with archaeal MC1 protein suggests gp275 as an architectural protein of Mimivirus involved in the condensation of genomic DNA. The expression profile of the R252 gene indicates its expression in the intermediate stage of infection with maximum expression at 6 h post-infection (p.i.) (Supplementary Fig. 1). The gene expression remained high at later stages of infection and comparable to the Mimivirus gene L425 (gene encoding capsid protein, gp455). The intermediate to late gene expression of Mimivirus R252 coincides with DNA replication and synthesis of mature capsids and, thus could work in tandem with other packaging proteins for DNA packaging.

### gp275 is present inside the Mimivirus capsid

Since all lineages of mimiviruses showed the presence of gp275 homologs, we carried out the mass spectrometry-based proteomics of purified APMV virions along with the Mimivirus isolates from our lab, Powai-lake megavirus, PLMV and Mimivirus Bombay, MVB^50,51^. The presence of gp275 in the APMV virion proteome and its homologs within the capsids of PLMV and MVB further strengthens their possible involvement in DNA organization (Fig. 2a). The packaging of gp275 within the capsids was also analyzed by imaging amoeba cells infected with the genetically modified Mimivirus. The genome of the Mimivirus was modified by inserting EGFP and mCherry genes in-frame, at the C-terminal end of R252 and L425 genes, respectively (Fig. 2b). This genetically modified Mimivirus, expressing green fluorescent protein labelled gp275 (gp275-EGFP) and red fluorescent protein labelled gp455 (gp455-RFP), was used for the co-localization study. At both 6 and 8 h post-infection (p.i.), gp275-EGFP and gp455-RFP localized in the viral factory (VF; site in the host cytoplasm, where viral replication, transcription, and assembly take place) along with DAPI staining (Fig. 2c, panel i). The co-localization of both the proteins with the viral DNA during infection and the observation of red and green fluorescence in the purified modified virus particles (Fig. 2c, panel ii) indicate the possible recruitment of gp275 for viral DNA arrangement during capsid morphogenesis and its subsequent packaging within the capsid.

**Fig. 2.**
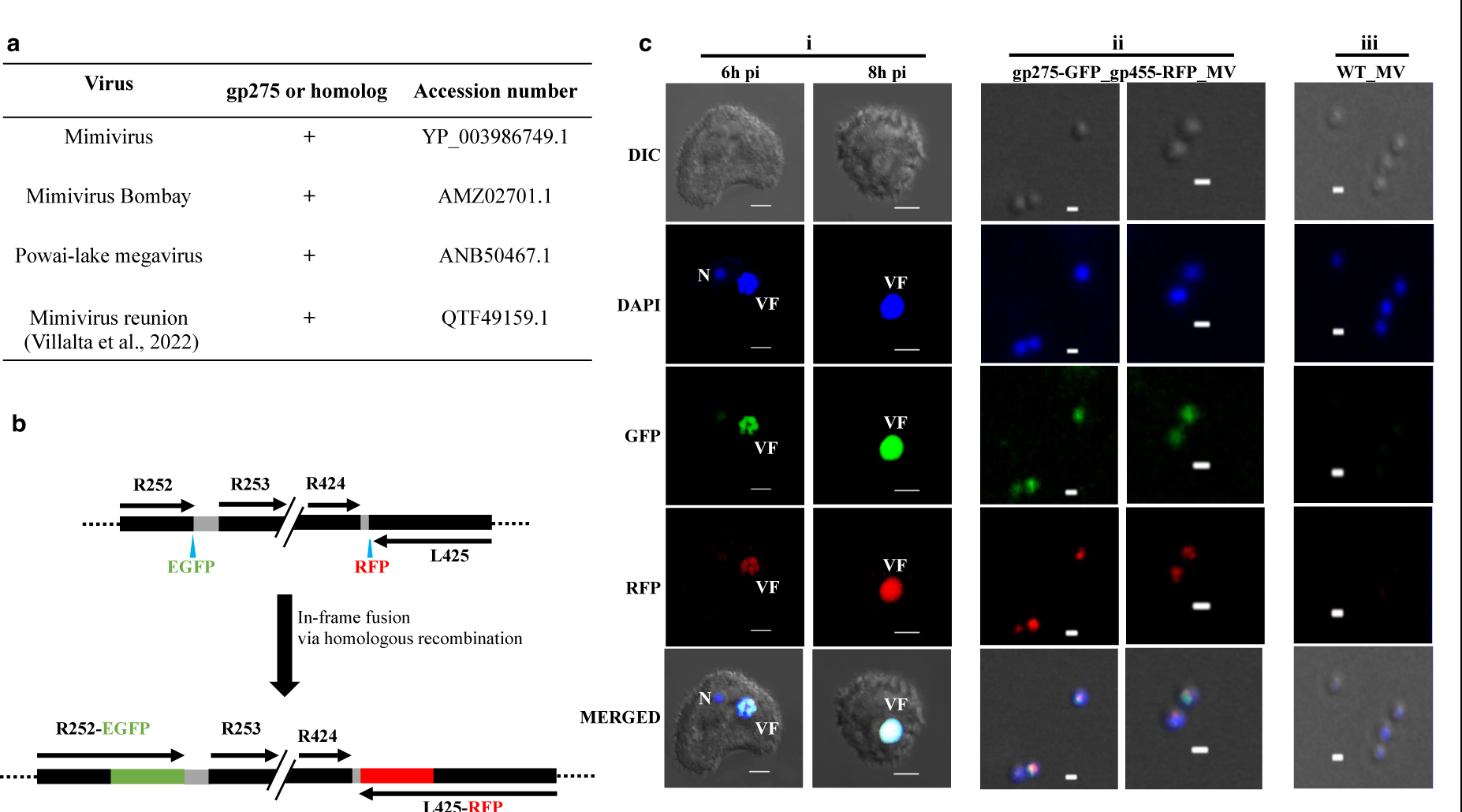
Mimivirus gp275 is packaged in the progeny virions. **a,** The putative gp275 homologs identified within the capsids of Mimivirus, Mimivirus Bombay, Powai-lake megavirus and Mimivirus reunion. **b,** Schematic showing homologous recombination strategy employed for the in-frame fusion of EGFP (enhanced green fluorescent protein) gene downstream to Mimivirus R252 gene in the virus genome for synthesising EGFP tagged gp275 protein. Similarly, gene L425 encoding capsid protein 1 (gp455) was tagged with RFP (red fluorescence protein) gene by in-frame fusion of mCherry to L425 gene in the virus genome. **c,** Laser scanning microscopy fluorescence images of *A. castellanii* cells infected with Mimivirus lysate expressing EGFP tagged gp275 and mCherry tagged gp455 at different time points, 6 and 8 h p.i. (panel i). Upon Mimivirus infection, gp275-EGFP and gp455-RFP co-localize with the viral DNA (shown by DAPI staining) in the VF. DAPI staining remains in the nucleus (N) throughout the infection, but at 8 h p.i., the intense fluorescence in the late VF hides the staining of the nucleus (Scale bar-5 µm). Panel ii is purified modified Mimivirus particles showing green fluorescence from R252-EGFP and red fluorescence from L425-RFP gene fusions (Scale bar-500 nm). Panel iii is the wild-type Mimivirus showing only DAPI staining (Scale bar-500 nm).

### gp275 induces DNA bending and condensation

To study the DNA binding properties of the putative MC1-like gp275 protein of Mimivirus, the corresponding R252 gene was cloned into the pET-28a plasmid. Additionally, based on the amino acid sequence comparison of Mimivirus gp275 with the archaeal MC1 protein, we constructed two more clones, namely, gp275-K126A, with Lys at position 126 mutated to Ala, and gp275-do, the minimal MC1 domain of gp275. gp275-do clone includes the structurally ordered part of gp275 with similarity to MC1 protein (residues K97-G228) excluding the disordered N and C-terminal regions (Fig. 3a).

**Fig. 3.**
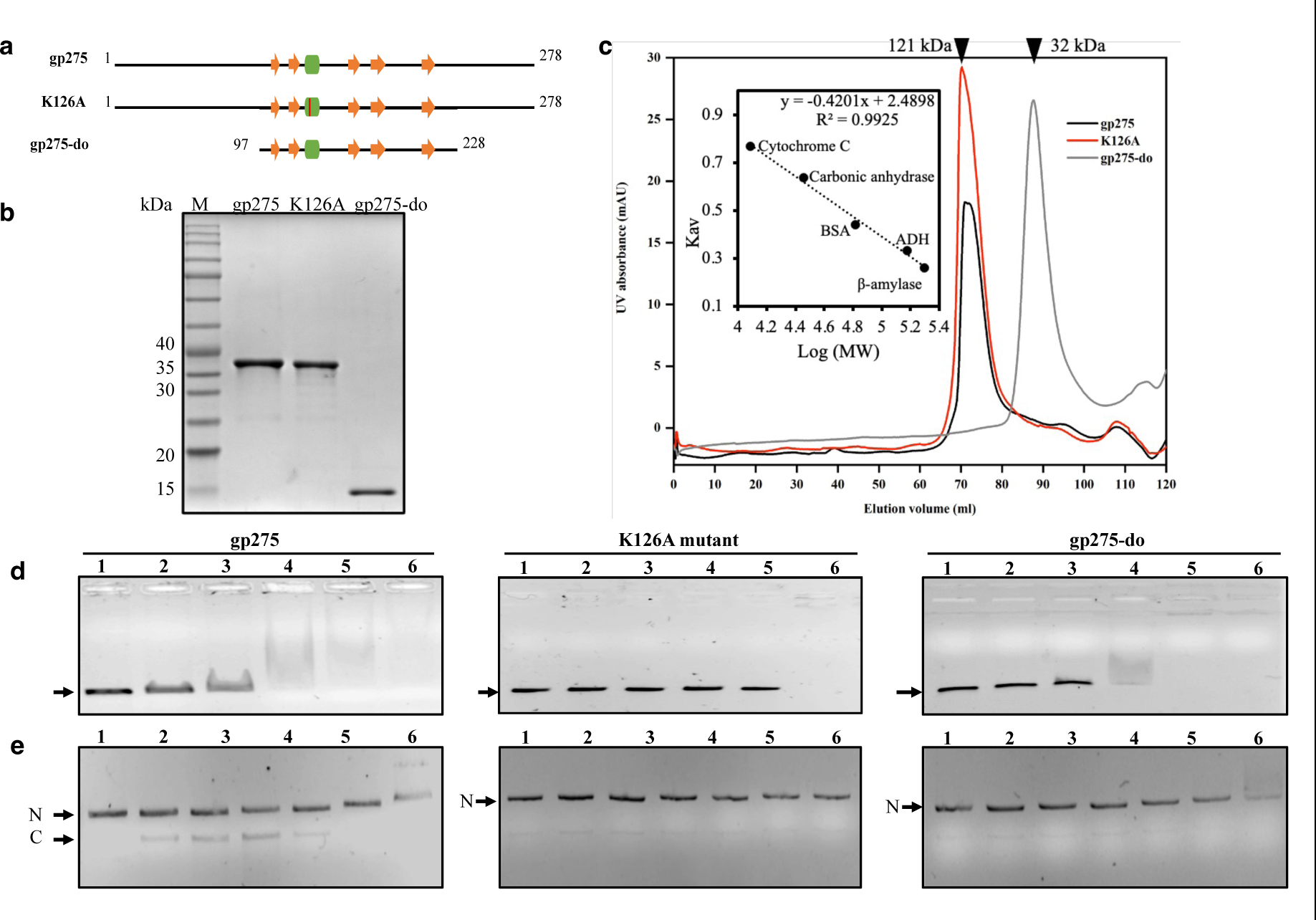
gp275 binds and supercoils DNA. **a,** MC1 domain organisation of the Mimivirus wild-type gp275, gp275-K126A mutant (red marks the mutation site) and gp275 minimal domain (gp275-do) depicting the position of β-sheets (orange) and ɑ-helix (green). The length of proteins and positioning of domain structural units are shown approximately to scale. The numbers at the start and the end depicts the amino acid position. **b,** Size exclusion chromatograms display elution profiles of proteins. For gp275 and gp275 mutant (K126A) elution peaks were observed at 71 ml elution volume corresponding to 121 kDa, which is predominantly tetrameric form. In the case of gp275-do, the elution peak at 88 ml corresponds to the dimeric form. Calibration chart for Hi-Load 16/600 Superdex-200 prep grade column shown as inset. **c,** Purified gp275, gp275 mutant (K126A) and gp275-do proteins analysis on Sodium Dodecyl Sulphate-Polyacrylamide (SDS-PA) gel. **d,** 1% agarose gel stained with ethidium bromide showing DNA band shift when 50 ng of 500 bp dsDNA was incubated with increasing concentration of proteins. The protein concentrations (nM) from lanes 1 to 6 were 0, 40, 80, 200, 400 and 600. The black arrow points at the unbound free DNA. **e,** Titration of the nicked pUC19 plasmid with proteins. The protein concentrations (nM), from lanes 1 to 6, were 0, 3, 6, 12, 25, 50, and 100, and the DNA concentration was 1 nM. Complexes were formed in 10 mM Tris-HCl, pH 7.5, 1 mM EDTA, and 50 mM NaCl. Electrophoresis was performed on a 1% agarose gel in 1X-TBE buffer. N-nicked plasmid, C-compacted plasmid. The faint bands observed below the nicked plasmid bands in the case of K126A and gp275-do are the contaminating supercoiled plasmid form in the substrate, nicked pUC19 DNA.

The expressed proteins were purified using metal affinity chromatography followed by size exclusion chromatography. Both gp275 and gp275-K126A eluted as predominant oligomeric form that would correspond to a tetramer of ∼121 kDa if assuming a standard protein density and globular quaternary structure for gp275 (independently confirmed by SEC-MALS analysis, data not shown). The MC1 minimal domain of gp275 (gp275-do) on the other hand, eluted as a dimer (∼ 32 kDa) under similar assumptions (Fig. 3b). All three purified Mimivirus proteins (Fig. 3c) were concentrated to ∼ 3 mg/ml and stored at −80°C. The DNA binding property of gp275, gp275-K126A and gp275-do was examined by the electrophoretic mobility shift assay (EMSA). The DNA-protein reaction mixtures were analyzed for binding on a 1% agarose gel. gp275 binding to DNA is evident by the shift seen in the DNA migration upon the addition of protein. With increasing protein-to-DNA ratios, a mixed population of varying DNA-protein complexes and subsequently complete shift close to the well was observed (Fig. 3d). gp275-do showed DNA binding comparable to gp275, while gp275-K126A exhibited reduced DNA binding as expected (Fig. 3d).

We also investigated the effect of protein binding on the conformational state of the relaxed circular DNA by performing the titration of nicked pUC19 DNA with increasing protein concentrations. In the presence of gp275, the appearance of a fast-moving species of DNA was observed, which is most likely due to the compaction/supercoiling of DNA. At higher concentrations of gp275 (25 nM and above), DNA migrated slower than the nicked DNA suggesting the formation of larger complexes with gp275 (Fig. 3e). The fast-migrating species was not observed in the case of both gp275-K126A and gp275-do proteins indicating the reduced ability of these proteins to compact the DNA.

The compaction of the nicked plasmid DNA by gp275 was further validated by examining gp275-induced DNA-bending using steady-state FRET (Fluorescence resonance energy transfer) analysis and compared with gp275-K126A mutant and the minimal domain, gp275-do. The fluorescence intensity of the fluorophore was used to monitor the change in distance between the 5’ ends of the duplex DNA labelled with 6-FAM (donor, fluorophore) and TAMRA (acceptor, quencher). The change in the distance between the 5’ ends of the duplex DNA labelled with 6-FAM (donor, fluorophore) and TAMRA (acceptor, quencher) was monitored by the variation in the fluorescence intensity of the fluorophore. Donor/acceptor labelled DNA of 70, 140, or 400 bp length, when incubated with Mimivirus gp275, demonstrated a decrease in the fluorescence intensity compared to the labelled DNA in the absence of gp275 protein. The reduction in the donor/acceptor intensity resulting from the change in the distance between both the 5’ ends of DNA indicates the bending of DNA by gp275 binding (Fig. 4a).

**Fig. 4.**
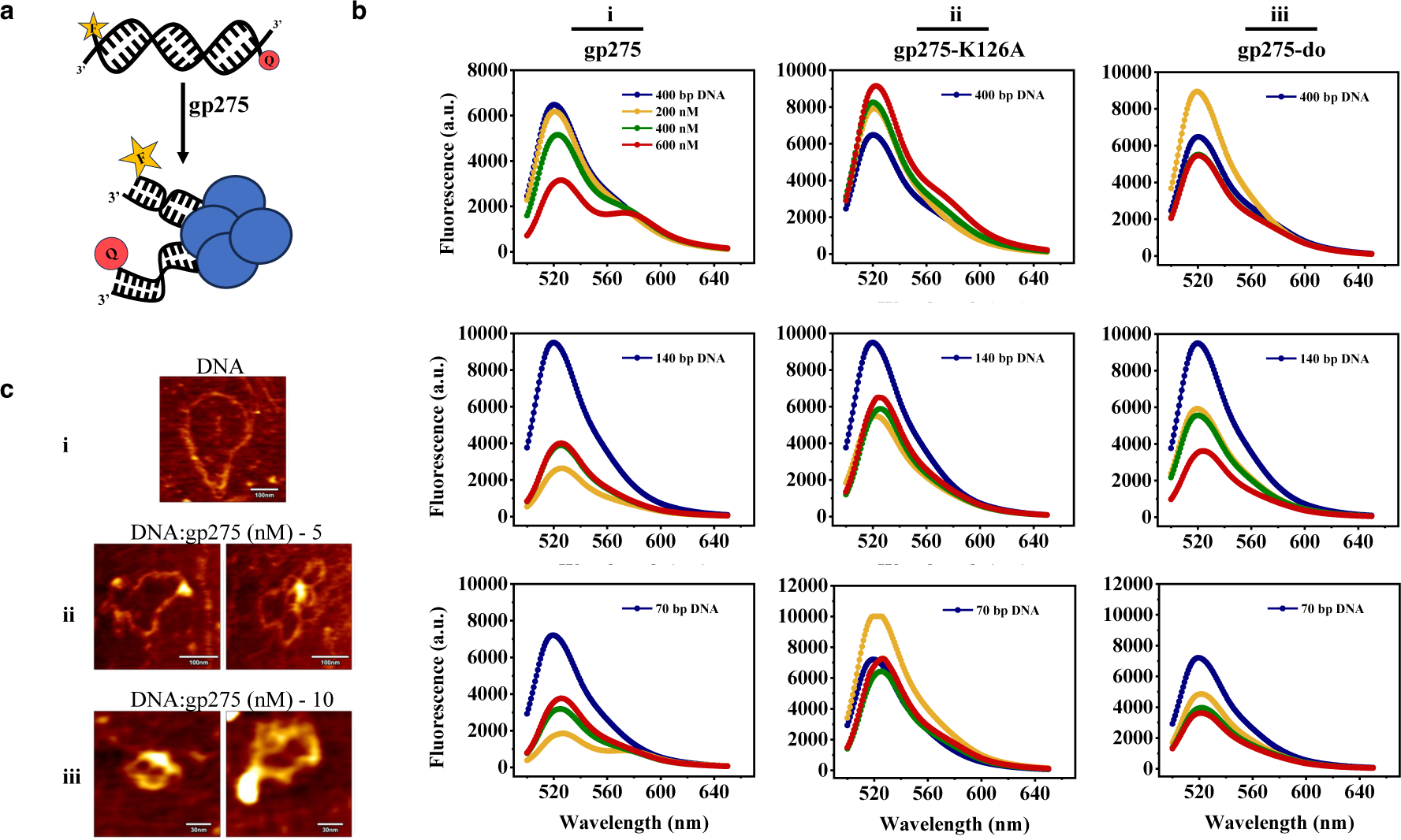
Binding of Mimivirus gp275 induced bending of the DNA substrate. **a,** Schematic illustration of labelled dsDNA substrate design with respective 5’ ends labelled with 6-FAM (F-Fluorophore) and TAMRA (Q-Quencher) and its bending upon gp275 (shown as tetramer, blue) binding. **b**, Emission spectra of labelled free and protein-bound DNA following excitation at 490 nm. The emission spectra are depicted from 500-650 nm for gp275, gp275-K126A and gp275-do binding to labelled DNAs (dark blue) at varying protein concentrations: 200 nM, yellow; 400 nM, green and 600 nM, red. **c**, Atomic force microscopy (AFM) imaging of nicked pUC19 plasmid DNA (2.6 kb) and DNA-gp275 complexes at DNA to protein molar ratio of 5 and 10. Scale bar in i, ii - 100 nm and iii – 30 nm.

The decrease in the intensity with increasing concentrations of gp275 is more evident with 70 bp and 140 bp DNA than with 400 bp DNA (Fig. 4b, panel i). In the case of gp275-K126A mutant, no decrease in the fluorescence intensity was observed with 400 bp and 70 bp labelled DNAs, whereas with 140 bp labelled DNA, some binding and bending was observed (Fig. 4b, panel ii). A decrease in fluorescence intensity was observed with gp275-do binding to the labelled DNAs. The decrease was greater with the shorter 70 bp DNA (Fig. 4b, panel iii). In agreement with the gel-based DNA bending, the binding of wild-type gp275 to the labelled DNA resulted in a decrease in fluorescence, indicating the bending of DNA upon protein binding. The gp275-K126A mutant was less efficient in inducing DNA bends implying lysine (K126) is a critical residue for DNA binding and bending. The Mimivirus wild-type gp275 was incubated with nicked circular pUC19 DNA and deposited on mica for atomic force microscopic (AFM) analysis. Free Nicked pUC19 DNA was observed in a relaxed circular form on mica (Fig. 4c, panel i). With increasing gp275 concentrations, progressive conformational changes in the relaxed DNA to a condensed state were observed (Fig. 4c, panels ii, iii).

### Mimivirus gp275 co-localizes with viral factories during infection

For establishing the localization of gp275 during Mimivirus infection, we performed fluorescence imaging of *A. castellanii* cells that were transfected with a recombinant plasmid (R252-pUbg) expressing the EGFP-tagged gp275. The infected amoeba cells were imaged at different time points post-infection and compared with uninfected amoeba for gp275 localization (Fig. 5). The untransfected (UT) cells without infection act as a control and do not show any EGFP fluorescence but a DAPI-stained host nucleus (Fig. 5, panel UT). In the transfected cells without Mimivirus infection (UI), gp275-EGFP remained dispersed in the cell cytoplasm but predominantly in the cell nucleus, suggesting the affinity of the protein towards DNA (Fig. 5, panel UI). The change in the pattern was observed when the *A. castellanii* cells transfected with R252-pUbg were infected with Mimivirus. At 4h p.i., gp275-EGFP was seen localized to both the cell nucleus and viral factory (Fig. 5, panel 4h pi). At 6h p.i., gp275-EGFP accumulated predominantly in the VF and by 8h p.i., gp275-EGFP was only visible in the VF (Fig. 5). The co-localization of the heterologously expressed Mimivirus gp275-EGFP with the newly synthesized viral DNA in the VF during infection indicates the possible recruitment of protein for its role in viral genome packaging. We also imaged *A. castellanii* cells transfected with pUbg-EGFP-R252 plasmid for modifying the Mimivirus genome by homologous recombination to endogenously express the green fluorescent-tagged gp275 protein (engp275-EGFP). The in-frame fusion of the EGFP tag to the R252 gene resulted in the expression of fluorescently labelled gp275 protein. There was no detection of green fluorescence in transfected but uninfected cells (Supplementary Fig. 2, P1-UI). Upon Mimivirus infection, green fluorescence appeared in the viral factory at 4h p.i. and a subsequent increase in the fluorescence signal was observed at later time points (Supplementary Fig. 2). The viral lysate (P1 generation), obtained after complete lysis was collected and used for subsequent amoeba cell infection to produce P2 generation. Green fluorescence was observed in the VF at both 6 and 8 h p.i. for P2 generation (Supplementary Fig. 2) confirming the endogenous expression of EGFP-tagged gp275.

**Fig. 5.**
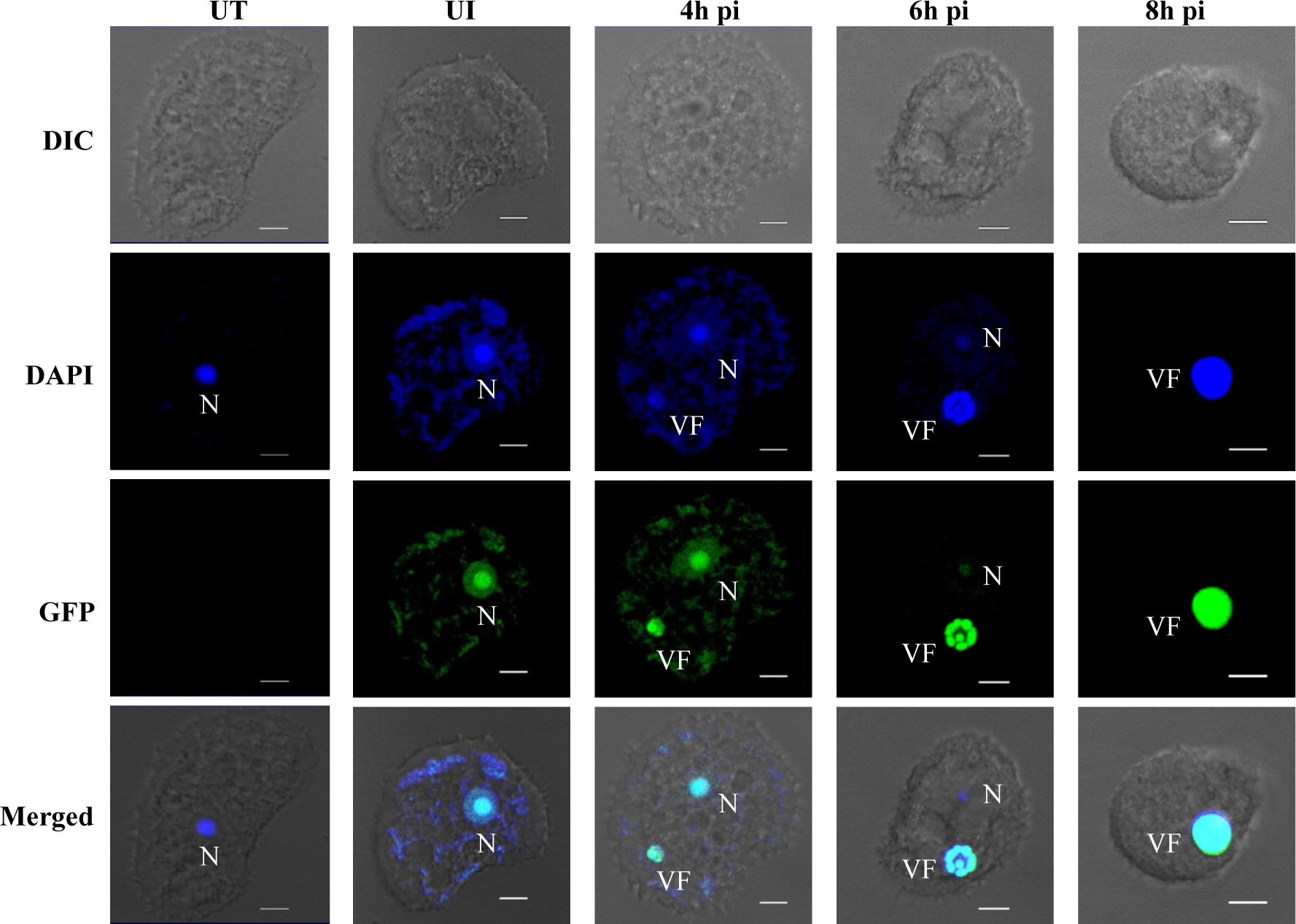
Mimivirus gp275 co-localized with the viral factory during infection. Laser scanning confocal microscopy fluorescence images of A. castellanii cells transfected with Mimivirus gp275-EGFP at different time points post-infection. In the uninfected (UI) cells, gp275-EGFP is scattered in the entire cell (including the nucleus, N). Upon Mimivirus infection, gp275-EGFP re-localises to the viral factory (VF). DAPI staining remains in the nucleus throughout the infection, but the intense fluorescence in the late VF hides the staining of the nucleus at 8 h p.i. In the case of untransfected (UT) cells, no green fluorescence was observed. The scale bar is 5 μm.

### Knockout of the R252 gene affects Mimivirus multiplication and infection

To understand the effect of R252 gene knockout (KO) on Mimivirus multiplication and infection, we employed homologous recombination for gene replacement. Homologous regions flanking upstream and downstream of the R252 gene were cloned into pUbg plasmid at the N and C-terminal of the EGFP gene, respectively. *Acanthamoeba castellanii* cells were transfected with linearized recombinant plasmid and further infected with Mimivirus. As shown in Fig. 6a, the homologous recombination results in the replacement of R252 with the EGFP gene in the viral genome leading to the formation of R252_KO-Mimivirus. The infection assessment, carried out by PCR (Supplementary Table 1), showed the presence of recombinant Mimivirus (R252_KO-Mimivirus) carrying EGFP gene in place of R252 after the first generation, P1 (Fig. 6b). Subsequent generations were monitored for the presence of R252_KO-Mimivirus by infecting amoeba cells with the viral lysate from the previous generation. The R252_KO-Mimivirus were outcompeted by the wild-type (WT) Mimivirus leading to the amplification of only WT-Mimivirus particles by the P4 generation (Fig. 6b). This illustrates that the R252 gene is essential for Mimivirus and its knockout has a significant effect on the production of viable virus particles for infection. To further analyze the effect of R252 knockout on Mimivirus multiplication, we measured the viral titer during infection by WT-Mimivirus and R252_KO-Mimivirus.

**Fig. 6.**
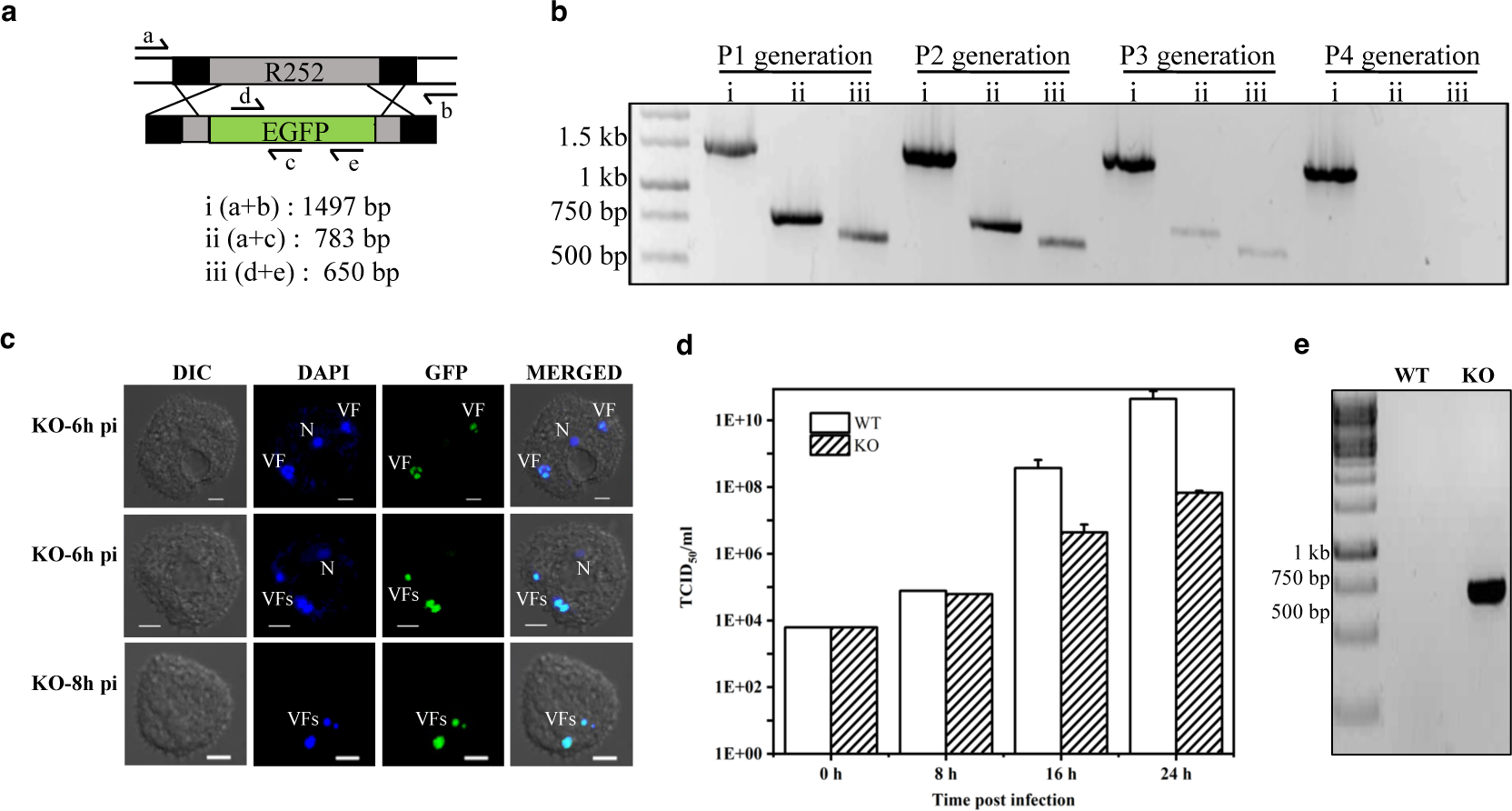
The R252 gene knockout affects the fitness of progeny virions. **a,** Schematic representation of the homologous recombination strategy used to generate the Mimivirus R252 gene knockout. Expected PCR product sizes upon recombination are also shown. Primer a: R252_KO_Forward; primer b: R252_KO_reverse; primer c: EGFP_KO_reverse; primer d: EGFP_internal_forward; primer e: EGFP_internal_reverse (supplementary table 1). **b,** 1% agarose gel shows PCR products demonstrating the integration of EGFP in the locus of Mimivirus R252. Homologous recombination observed as, a+b: wild type locus; a+c: 5’ integration; d+e: EGFP gene. **c,** Laser scanning confocal microscopy-based fluorescence imaging of *A. castellanii* cells infected with R252_KO-Mimivirus at different time points p.i. Infected amoeba cells displayed a highly stained nucleus (N) and scattered VFs at 6 h p.i. (top two panels) and smaller (3-4 in number) atypical viral factories (VFs) at 8 h p.i. Scale bar-5µm. **d,** Based on TCID50/ml calculation, the bar graph demonstrates Mimivirus multiplication. Titres were determined for WT-Mimivirus and R252_KO-Mimivirus at the designated time points (0, 8, 16 and 24 h p.i.) and values represent the mean of results from three independent experiments. **e,** PCR amplification of EGFP gene (657 bp) from R252_KO-Mimivirus lysate.

We also imaged *A. castellanii* cells transfected with recombinant plasmid upon Mimivirus infection using confocal microscopy. With the generation of R252_KO-Mimivirus upon recombination during viral replication, we observed dense DAPI staining in the nucleus till 6 h p.i. (Fig. 6c, panel KO-6h pi) which stains faintly in the case of WT-Mimivirus infected amoebal cells. In addition to this, in most of the transfected cells, small isolated VFs were observed at 8 h p.i. (Fig. 6c, panel KO-8h pi) whereas a single large coalesced VF is predominantly observed after 8 h in the case of WT-Mimivirus infected amoebal cells (Fig. 5). By using the end point dilution assay, the difference in the virus production rate was estimated in samples collected at 0, 8, 16 and 24 h p.i. Significant reduction in the viral titer in the case of R252_KO-Mimivirus infection was apparent, particularly at later stages of infection i.e. 16 h and 24 h p.i. (Fig. 6d). Thus, the knockout of the R252 gene affects Mimivirus multiplication and leads to a significant reduction in the viral titer. The genomic DNA was isolated from the viral lysate obtained at 24 h p.i. and PCR amplification was carried out to confirm the presence of the EGFP gene in the R252_KO-Mimivirus particles (Fig. 6e). This demonstrates that the lack of gp275 has a notable impact on the production of viable progeny virus particles, resulting in the formation of small aberrant viral factories in R252_KO-Mimivirus, which do not coalesce as observed in the case of WT-Mimivirus.

## Discussion

In this report, we show that Mimivirus codes for a homolog of the DNA-condensing MC1 protein (gp275) and demonstrate its presence in the Mimivirus virions by mass spectrometry and microscopy. Knocking out of the R252 gene (that codes for gp275) affects viral factory maturation and also results in the reduced viability of the virions assembled in the absence of gp275. The biochemical characterization and atomic force microscopy studies further suggest that the purified gp275 protein is a tetramer that binds, bends, supercoils, and condenses the DNA. This is the first report of a virus using an archaeal non-histone MC1-like protein for DNA compaction, adding to the diverse strategies adopted by viruses for condensing the packaged DNA inside the virion.

In the previous studies aimed at gaining insights into the Mimivirus genome organization, the compact arrangement of DNA in the Mimivirus particles has been described^33,34^. The huge genomes of large viruses with numerous packaged proteins necessitate the condensation of the genome and is an essential step in the viral life cycle^52,53^. The viruses belonging to the *Nucleocytoviricota* phylum are believed to condense the genome inside the capsid for packaging either by encoded histone proteins or non-histone proteins (Supplementary Table 2). The presence of one or more histone genes has been identified for viruses of the phylum *Nucleocytoviricota* such as marseilleviruses, pandoraviruses, iridoviruses, and medusaviruses. Iridoviruses and Medusavirus histones are detected in the capsids suggesting a role in DNA condensation though the formation of higher-order chromatin structure is not known^19,20^. Marseillevirus packages histone proteins that have been shown to form eukaryotic nucleosome-like condensates^18,36,54^. By contrast, the single histone protein H2B encoded by pandoraviruses is not detected in the virions inferring their role in suppressing host multiplication thus favouring viral replication^10, 55,56^. In the case of phycodnaviruses, the histone H3 variant gene was identified in the genome of Dishui lake phycodnavirus 1 (DSLPV1)^57^ and DNA-binding proteins called dinoflagellate/viral nucleoproteins (DVNPs) are encoded by most phycodnaviruses and are expected to be taking part in genome packaging ^58,59^. DNA condensing non-histone basic proteins (P64) are packaged in ascoviruses that sequester the genome for encapsidation and bacteria-like condensing protein, pA104R, in ASFV associates closely with the viral DNA^60,61,62^. In poxviruses, packaging proteins like A32 and I6 are essential for the translocation and packaging of DNA into the virion. Polyamines are identified in Vaccinia virions that might be involved in compacting DNA^79^. Thus, the molecular mechanisms of genome organization/condensation evolved by viruses of the *Nucleocytoviricota* phylum require diverse types of proteins that apparently are phylogenetically unrelated.

The 1.2 Mb genome of Mimivirus encapsidated in a ∼ 500 nm capsid shell does not encode any histone proteins. Still, two Mimivirus-coded GMC-oxidoreductases have been implicated as the major constituents of the proteinaceous shell of genomic fibers^34^. In the Mimivirus genomic fiber of ∼30 nm diameter, DNA is folded as five or six strands in its highly compacted conformation. However, studies have shown that both GMC-oxidoreductases are not essential for the survival of Mimivirus^35,36^ bringing into question which other protein/s is/are involved in DNA condensation. Our domain search and sequence alignment identified the putative MC1 domain in the Mimivirus gp275, encoded by gene R252 (Fig. 1a). The MC1 domain of archaea is involved in the genome organization^63^. It is structurally distinct from the other DNA-condensing domains such as the histone-fold found in eukaryotic and archaeal histones^42,45,64^. MC1 belongs to the architectural non-histone proteins and is found in archaea where Alba, the highly represented non-histone archaeal protein, is absent^7,65^. Archaeal MC1 is a relatively small basic protein that induces sharp turns in the DNA by neutralizing the negative charges of the proximal phosphates in the DNA with the side chains of critical arginine residues^39,41,43^. Mutation of a key lysine (K126) residue in gp275, equivalent to the conserved Arg residue of MC1, resulted in a much-reduced DNA binding affinity and DNA bending (Figs. 3 and 4). Interestingly, our sequence similarity search showed that the presence of gp275 homologs containing the MC1 domain is restricted to phycodnaviruses, apart from a single hit from the metagenome-assembled Marseillevirus (Fig. 1a).

The acquisition of MC1-domaining protein by Mimivirus could be attributed to lateral gene transfer from the host to virus^66^ (Fig. 1c). Phycodnaviruses and mimiviruses that carry MC1 homologues are also phylogenetically most closely related within the phylum *Nucleocytoviricota*^22,67^, and in addition, marine mimiviruses have an association with large algal viruses^68^. The identification of MC1 domain-containing proteins further strengthens the association between the two viral groups (Fig. 1).

Our virion proteome analysis detected gp275 inside the prototype Mimivirus capsid as well as in the other closely related Mimivirus virions isolated from the Indian subcontinent^50,51^ and the accumulation of EGFP-tagged gp275 within the viral factory during viral infection suggests the recruitment and involvement of gp275 in DNA condensation and packaging (Fig. 5). Additionally, *in vivo* studies confirm the importance of gp275 for Mimivirus multiplication and generation of viable particles (Fig. 6). Based on our observations, we speculate the potential involvement of gp275 in inducing bends in Mimivirus DNA and thus aiding the folding of the viral genome to be arranged as the genomic fiber inside the capsid in conjunction with the other packaging proteins^34,53^ (Fig. 7).

**Fig. 7.**
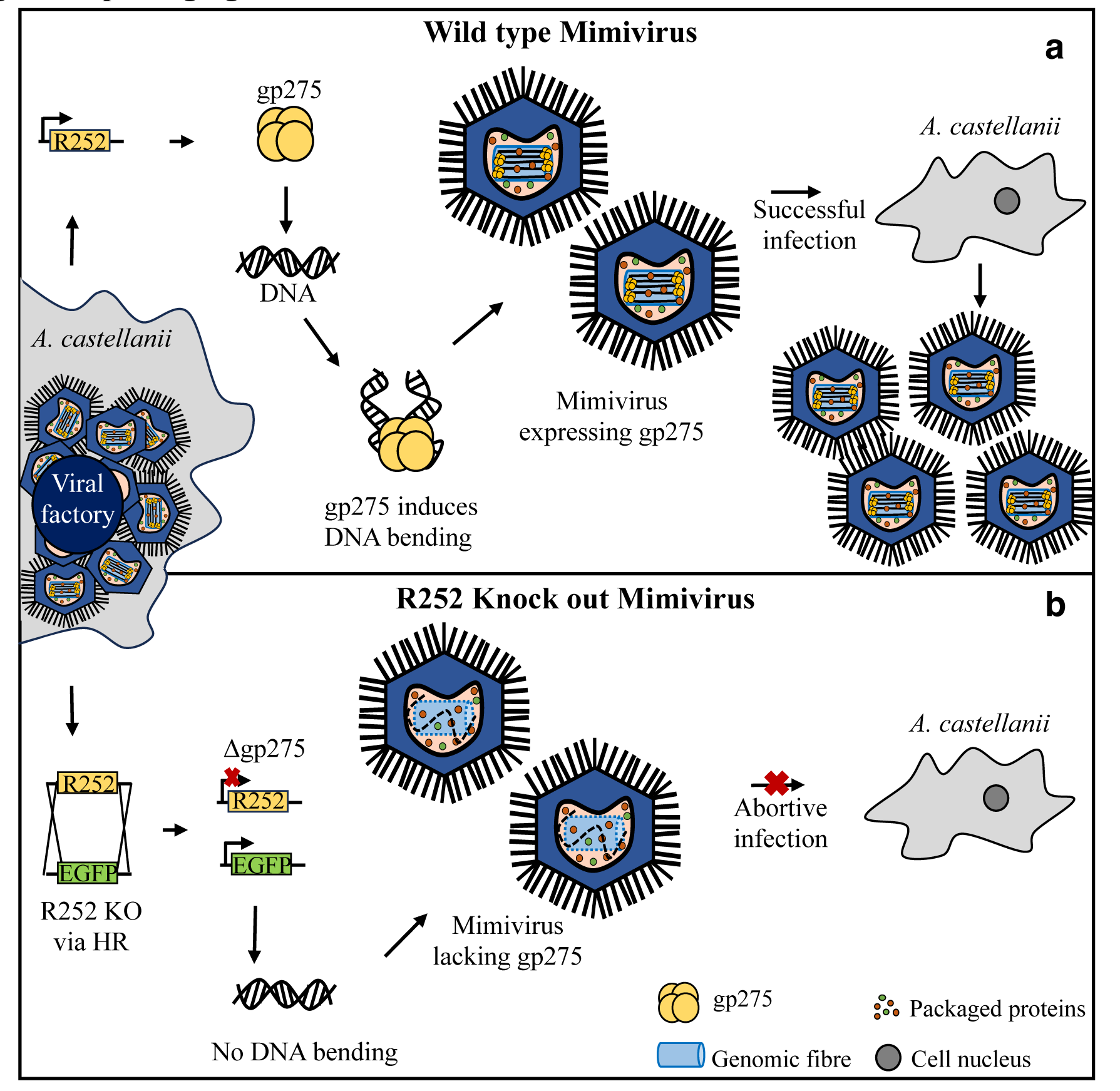
Schematic representation of Mimivirus gp275 role in DNA-bending and genome packaging. **a,** *Acanthamoeba castellanii* cell infected with Mimivirus expressing wild-type gp275. gp275 (yellow) induces bends in the DNA and facilitates genome packaging by folding DNA (black) in the genomic fiber (blue cylinder) leading to the generation of viable viral particles capable of successful infection. **b,** *A. castellanii* cells transfected with EGFP gene segment for R252 gene knockout (KO). Upon Mimivirus infection R252 is replaced with EGFP gene by homologous recombination (HR) leading to deletion of gp275 in the genome (Δgp275). Mimivirus lacking gp275 expression results in abortive infection. The generation of non-infectious viral particles in R252 knock-out Mimivirus could be either due to the lack of DNA packaging (shown as a dashed black line) inside the capsid or the absence of the formation of genomic fiber (dashed blue cylinder). Other packaged proteins are depicted in orange and green circles while *A. castellanii* nucleus in a grey circle.

With the findings of the present study, we propose gp275 as an architectural protein of Mimivirus involved in genome compaction and packaging. The organization of Mimivirus DNA in a ∼30 nm genomic fiber is speculated to occur either inside the capsid as the DNA is pumped by the putative packaging ATPase (gp468 coded by L437 in the case of Mimivirus^69^) through the 20 nm capsid portal^70^ or with the involvement of multiple packaging ATPase rings pumping the folded genomic fiber^71^. It is not clear whether, in the absence of gp275, there is no formation of genomic fiber and translocation of the Mimivirus genome into the preformed capsid (if considered the case of the folded DNA translocation) leading to the production of non-infectious mimiviruses. Or the genome is translocated into the capsid (if considered the case of DNA folding post translocation) but in the lack of gp275 expression, genomic DNA fails to fold and get accommodated into the genomic fiber along with the RNA polymerase subunits^34^ inhibiting early gene transcription upon unwinding following infection. The mechanism of DNA condensation by gp275 is yet to be understood, but can be speculated to work in a manner similar to archaeal genome condensation by MC1 protein which induces sharp bends in the DNA (Loth et al., 2019). The studies focussing on deducing the structure of gp275-bound DNA would help in gaining insights into the mechanism employed by gp275 for DNA condensation.

## Methods

### gp275 sequence analysis and structure prediction

Mimivirus gp275 amino acid sequence was retrieved from NCBI and used as a query to search for homologous sequences. The amino acid sequence similarity search was performed using the PSI-BLAST with default settings. The multiple sequence alignment of the homologous sequences was performed using Clustal Omega^46^ with default settings. The gp275 amino acid sequence was subjected to online tools such as InterPro Scan and Pfam to identify the protein domain. Phylogenetic tree construction of Mimivirus gp275 with homologous sequences from other viruses and MC1 protein sequences of archaea was done using the MEGA7 program^73^. A 3D structural model for gp275 was generated using AlphaFold2^74^ and visualized using PyMOL^72^. A total of 5 models were created and the top-ranked model with the best pLDDT score is represented^44,74^.

### *Acanthamoeba castellanii* culture and Mimivirus isolation

*A. castellanii* cells were cultured at 28°C in PYG medium (2% proteose peptone, 0.1% yeast extract, 4 mM MgSO_4_, 0.4 mM CaCl_2_, 3.4 mM sodium citrate, 0.05 mM Fe(NH_4_)_2_(SO_4_)_2_, 2.5 mM Na_2_HPO_4_, 2.5 mM KH_2_PO_4_ pH 6.5, 100 mM glucose) using T-25 flask. For Mimivirus isolation, amoeba cells at a density of 10^5^ cells/ml in 4 ml culture medium were seeded in a T-25 flask and cultured overnight at 28°C. After 80% confluency was reached, the cells were infected with Mimivirus at an MOI of 10 and incubated at 32°C. On completion of cell lysis, the lysate was centrifuged at 800 *g* for 15 min to remove the cell debris. The supernatant obtained was then centrifuged again at 15,000 *g* for 90 min to pellet down the viruses. The viral pellet obtained was resuspended and stored in sterile phosphate-buffered saline (PBS). The viral titre was quantified by calculating the median tissue culture infectious dose (TCID_50_)^75^.

### Mass spectrometry

Powai Lake Megavirus (PLMV) and Mimivirus Bombay (MVB) were isolated previously by our lab using the same protocol as APMV^50,51^. Viruses were purified using 30% to 60% sucrose density gradient centrifugation at 5,000 *g* for 20 min and washed with PBS. Purified virus preparations were analysed by SDS-gel electrophoresis and corresponding bands were subjected to in-gel trypsin digestion. The treated peptides were subjected to a Q Exactive Plus Orbitrap mass spectrometer (Thermofisher Scientific) instrument. The data obtained was analysed using the MaxQuant and/ or the Proteome Discoverer software.

### Cloning and purification of gp275, gp275-domain and gp275-K126A

Mimivirus R252 gene coding for gp275 protein was amplified from the APMV genome by polymerase chain reaction (PCR) using the primers, (Forward: 5’-CTAGCTAGCATGTCAACTCGTTCCAAC −3’ and Reverse: 5’-CTCGGATCCTTAACGACTAGCTTTAGC −3’). The amplified gene was cloned into *NheI* and *BamHI* restriction sites of the pET28a vector to yield an N-terminal polyhistidine-tagged protein. The K126A mutant, with alanine replacing lysine at the 126^th^ position in the putative MC1 domain of gp275, was created by site-directed mutagenesis using the primers (Forward: 5’- GGTGGTACTCCTGCArCAAGCTG −3’ and Reverse: 5’-CAGCTTGTGCAGGAGTACCACC −3’) and cloned into *NheI* and *BamHI* restriction sites of the pET28a vector. To clone the putative MC1 domain of gp275, the domain sequence was amplified using primers (Forward: 5’- TCTCCATGGGGAAAGGTGATGAATCTC −3’ and Reverse: 5’- TTCAAGCTTTGACTTCTTGTTGGATCGG −3’) and inserted into *NcoI* and *HindIII* restriction sites of the pET28a vector. The recombinant plasmids were then utilized to transform *E. coli* BL21 (DE3) RIPL cells for protein overexpression. Large-scale cultures were grown in a Luria-Bertani (LB) medium containing 25 μg/ml chloramphenicol and 50 μg/ml kanamycin until an optical density of 0.5-0.6 at 600 nm was reached. The cells were induced with 0.3 mM isopropyl-1-thio-β-d-galactopyranoside (IPTG) and allowed to incubate at 16°C for 15 h with constant shaking. The bacterial cells were harvested by centrifugation at 7,000 *g* and re-suspended in buffer A (50 mM Tris pH 7.4, 5 mM imidazole, 1 M NaCl, 5% glycerol, 1 mM Phenylmethylsulfonyl fluoride (PMSF) and 1 mM benzamidine hydrochloride). The cell lysis was performed using sonication at 40% amplitude for 20 min and the cell lysate was cleared using centrifugation at 14,000 *g* for 30 min. The soluble extract was loaded onto the Hi-Trap Chelating HP 1 ml pre-packed column (GE Healthcare/Cytiva) pre-equilibrated with buffer A containing 500 mM NaCl. Elution of gp275 was carried out with a 60-600 mM imidazole linear gradient in buffer A containing 500 mM NaCl. The protein fraction obtained was further subjected to Hi-Load Superdex 200 pg 16/600 size-exclusion chromatography column (GE Healthcare/Cytiva) equilibrated with buffer B (20 mM Tris pH 7.5, 100 mM NaCl and 5% glycerol). The purified protein fractions were concentrated to ∼3 mg/ml in buffer B using Amicon® Ultra 15 ml centrifugal filters and were stored in aliquots at −80°C. The protein concentration was determined by bovine serum albumin (BSA) standard curve using the BioRad dye reagent.

### Electrophoretic mobility shift assay

Electrophoretic mobility shift assay (EMSA) was carried out to detect DNA-protein interactions for all three purified proteins, gp275, gp275-K126A and gp275-do. The DNA template of 500 bp was generated by PCR from the pET28a-R252 plasmid. The binding buffer contained 20 mM Tris-Cl pH-8, 100 mM NaCl and 5 mM MgCl_2_. 50 ng of DNA was incubated with an increasing concentration of purified proteins for 1 h at 32°C. The whole reaction mix of 10 ml was then loaded on 1% agarose gel containing 1X TBE buffer and visualized using a UV-transilluminator (UVI-TEC).

### Supercoiling assay

A supercoiling assay was performed to observe the conformational changes in DNA upon purified protein binding. The nicking of duplex circular DNA is known to relax the plasmid with ethidium bromide (EtBr)-inhibited restriction digestion^76^. Nicked plasmid pUC19 DNA was prepared by *EcoRI* digestion in the presence of 150 µg/ml EtBr^77^ and purified using the GeneJET Gel Extraction Kit (Thermofisher Scientific) to be used as the substrate for supercoiling assays. 1 nM of nicked pUC19 DNA was complexed with varying concentrations of proteins, gp275, K126A and gp275-do. The reaction was carried out for 1 h at room temperature in the presence of 10 mM Tris pH 8, 1 mM EDTA, and 5 mM MgCl_2_. The complexes were analyzed on 1% agarose gel in 1X TBE buffer by EtBr staining.

### Fluorescence Spectroscopy

Fluorescence spectroscopy was employed to determine DNA bending upon protein binding. Duplex DNA of 70, 140 and 400 bp with 5’ ends labelled with 6-carboxyfluorescein (6-FAM) and tetramethylrhodamine (TAMRA) were amplified using primers (synthesized from Sigma-Aldrich, supplementary table 1) and incubated with gp275, gp275-K126A and gp275-do. The excitation wavelength used was 490 nm and fluorescence was measured from 500 to 650 nm.

### Atomic force microscopy

Purified gp275 was incubated with 0.2 nM nicked plasmid DNA at a DNA-to-protein molar ratio of 5 and 10 for 1 h at 37°C. Samples were diluted in 20 mM Tris-HCl pH 7.5 and 5 mM MgCl_2_. A 20 µl sample was deposited on the freshly cleaved mica sheet for 2 min, rinsed with 2 ml of water and dried. Atomic Force Microscopy (AFM) imaging was done using Asylum Research MFP-3D using a probe with a silicon tip of 70 kHz resonance frequency.

### Cloning of Mimivirus R252 gene for EGFP tagging

The expression vector, pUbg (kind gift from Hyun-Hee Kong, Dong-a University, South Korea), with the *Acanthamoeba* ubiquitin promoter and enhanced green fluorescent protein (EGFP) as the reporter gene was used to clone Mimivirus R252. The gene amplification was carried out using Mimivirus genomic DNA as a template with primers (Forward: 5’- CTTCCATGGATGTCAACTCGTTCCAAC −3’ and Reverse: 5’-CCGACTAGTTTAACGACTAGCTTTAGC −3’). The amplified gene was cloned into *NcoI* and *SpeI* restriction sites of the pUbg vector to generate a recombinant vector (pUbg-R252) expressing EGFP-tagged gp275.

### Fluorescence localization of gp275 in infected *A. castellanii* cells

1×10^6^ *A. castellanii* cells were transfected with 1 μg of pUbg-R252 plasmid using Effectene transfection reagent (QIAGEN) in a 60 mm dish following the manufacturer’s protocol. Transfection was carried out for 4 h at 28°C and then cells were washed with 1X PBS. Transfected *A. castellanii* cells were grown on poly-L-lysine (0.01%) coated coverslips in a 12-well plate and infected with Mimivirus at an MOI of 10 except for the negative control (amoeba cells only). At 4, 6 and 8 h post-infection (p.i.), cells were fixed with 1X PBS containing 4% formaldehyde for 20 min. After washing with 1X PBS, fixed cells were permeabilized with 0.2% Triton-X, followed by staining with 5 µg/ml DAPI (4′,6-diamidino-2-phenylindole) for 20 min. The coverslips were washed again with 1X PBS and mounted on a glass slide with a mounting medium and further observed for DIC, DAPI, and GFP fluorescence recording using a Carl Zeiss LSM 780 microscope (Zeiss, Germany).

### Gene knockout methodology and analysis

R252 gene knockout was carried out using homologous recombination for gene replacement. 550 bp of the homologous region upstream to the R252 gene was cloned into the pUbg vector at *NcoI* and *SpeI* sites using primers (Forward: 5’- TCTCCATGGATGTTTTTCGAAACCGAT −3’ and Reverse: 5’-TCCACTAGTCTCCAGATTGTTTAGTAG −3’). Similarly, 550 bp of the homologous region downstream to the R252 gene was cloned into the pUbg vector at the *XbaI* site using primers (Forward: 5’- TTCTCTAGACAAGCTGCCAGTAAAGGA −3’ and Reverse: 5’-CTCTCTAGAGGTTTGGACATGAGTGTA −3’). The orientation at the XbaI site was checked using flanking region primers (Forward: 5’- TCTCCATGGATGTTTTTCGAAACCGAT −3’ and Reverse: 5’-CTCTCTAGAGGTTTGGACATGAGTGTA −3’). The recombinant plasmid generated was digested with *SacI* and *NcoI* and used to transfect *A. castellanii* cells at a concentration of 1 μg using Effectene transfection reagent (QIAGEN). Cells were infected with Mimivirus (MOI-10) 4 h post-transfection and after 1 h p.i., the supernatant was removed to eliminate excess viral particles and fresh PYG was added to the cells. On completion of lysis, the viral pellet was collected and used for subsequent rounds of infection. After each passage, genomic DNA was isolated (Raoult et al., 2004) and PCR was performed to check for gene knockout. To estimate the effect of R252 gene knockout on virus multiplication, samples at 0, 8, 16 and 24 h p.i. were collected and viral titre at each time point was quantified using TCID_50_ calculation.

### Fluorescence gene tagging protocol

For the construction of a modified Mimivirus genome expressing tagged proteins, we utilized the recombination method for in-frame insertion of fluorescent tags like EGFP or mCherry at the C-terminal end of the target gene. To express green fluorescent tagged gp275, 720 bp of EGFP gene was amplified from pUbg plasmid using primers (Forward: 5’- AGCTGGCGCAAAAAAAGCACCAGCTGCTAAAGCTAGTCGTATGGTGAGCAAGGGCGAGGA −3’ and Reverse: 5’- ATATATTATAACTACATAATTTTTATACAAATCAAATTGTTACTTGTACAGCTCGTCCATGCC −3’). Similarly, for expressing red fluorescently tagged Mimivirus capsid protein (gp455), 711 bp of mCherry gene was amplified from pAW8-mCherry plasmid using primers (Forward: 5’- CAGAATGCTTAGCGGAATGGCCGGATTAGCGTACAGTAATATGGTGAGCAAGGGCGAGGA −3’ and Reverse: 5’- CAGAATGCTTAGCGGAATGGCCGGATTAGCGTACAGTAATATGGTGAGCAAGGGCGAGGA −3’). The amplified products were gel-purified and used for transfecting amoeba cells with the Effectene transfection reagent (QIAGEN). The transfected amoeba cells were subjected to infection with wild-type Mimivirus at an MOI of 10 and incubated till complete cell lysis. The cell debris was removed by centrifugation at 800 *g* for 10 min. The supernatant obtained was centrifuged at 15,000 *g* for 90 min at 4°C and the virus pellet obtained was dissolved in Millie Q and stored at 4°C. These Mimivirus particles with modified genomes were used to infect amoeba cells and observed using confocal microscopy for DIC, DAPI, GFP or RFP. For endogenous EGFP-tagged gp275 expression, two subsequent generations (P1 and P2) of Mimivirus infection were imaged.

## Acknowledgement

This research in KK lab is presently funded by the Board of Research in Nuclear Sciences, BRNS, India [58/14/11/2020-BRNS/37188], the Department of Biotechnology, DBT [BT/PR35928/BRB/10/1841/2019], the Science and Engineering Research Board [CRG/2023/00131].

