## Supplemental for "Mimivirus encodes an essential MC1-like non-histone architectural protein involved in DNA condensation"

Deepti Sharma, Fasseli Coulibaly and Kiran Kondabagil*

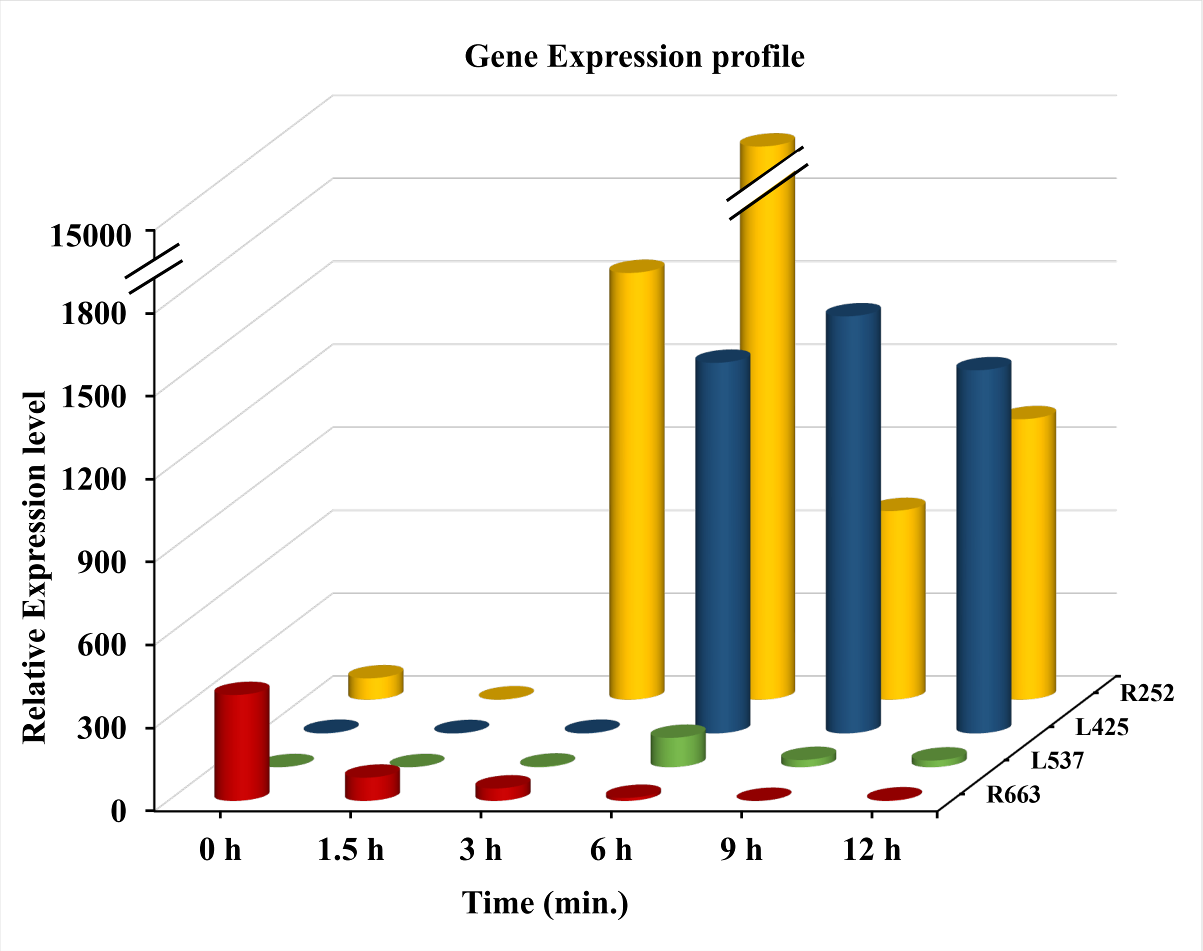


**Supplementary Figure 1 Expression profile of Mimivirus R252.** Relative expression level of R252 along with early (R663), intermediate (L537) and late (L425) mimivirus genes showing intermediate to late expression of R252 gene. Data procured from http://www.igs.cnrs-mrs.fr/mimivirus/ and the transcriptome analysis details can be found in Legendre et al., 2010.

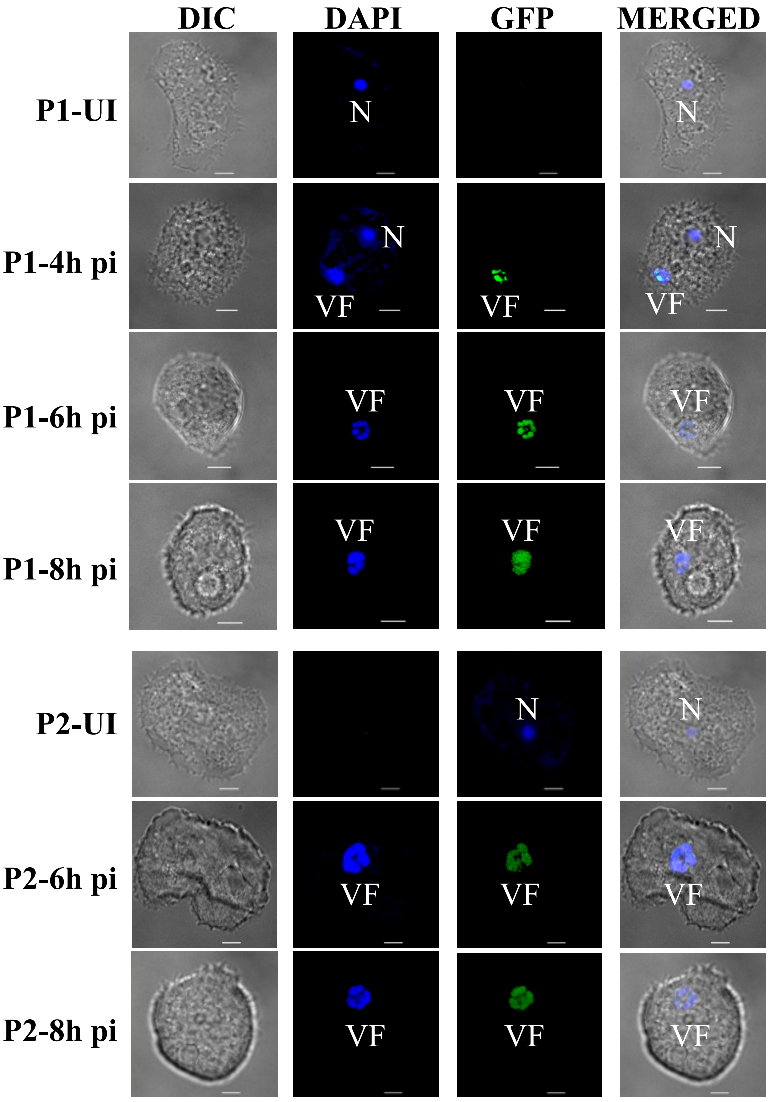


**Supplementary Figure 2: Mimivirus endogenously expressed gp275-EGFP co-localization.**

Laser scanning confocal microscopy-based fluorescence imaging of transfected *A. castellanii* cells expressing endogenous EGFP tagged gp275 (engp275-EGFP) at different time points post Mimivirus infection. Expressed engp275-EGFP co-localises with the viral factory (VF) for two subsequent generations (P1 and P2). DAPI staining was observed in both the nucleus (N) and viral factory (VF). Scale bar- 5µm.

**Supplementary Table 1: Oligonucleotides used in the present study.**

| Sr. No. | Sequence (5'-3) / primer name | Application |
|---|---|---|
| 1. | CTGATTCGTTTTCATTTGGAGT / a | Diagnostic PCR for R252 gene KO |
| 2. | GTAACGGAACTCCCTCTGAT / b | Diagnostic PCR for R252 gene KO |
| 3. | GAACTTGTGGCCGTTTACGTC / c | Diagnostic PCR for R252 gene KO |
| 4. | GACGTAAACGGCCACAAGTTC / d | Diagnostic PCR for R252 gene KO |
| 5. | CTTGTACAGCTCGTCCATGC / e | Diagnostic PCR for R252 gene KO |
| 6. | FAM-CAATCAGGTGGTAAAGGTGATG | 5'-FAM labelled primer |
| 7. | TAMRA-ATCTACCAAATGACCTTCCAGT | 70-mer labelled oligo |
| 8 | TAMRA-TCAGCTTTAATTTTGTGAATCA | 140-mer labelled oligo |
| 9. | TAMRA-TCTCCTGACTTCTTGTTGGATC | 400-mer labelled oligo |

**Supplementary Table 2: Histone and non-histone proteins encoded by NCLDVs.**

| NCLDV family | Example | Histone proteins | Non-histone proteins |
|---|---|---|---|
| *Ascoviridae* | *Spodoptera frugiperda* ascovirus | ND | P64 |
| *Asfarviridae* | African swine fever virus | ND | pA104R |
| *Iridoviridae* | Invertebrate iridiovirus 9 | H4, H3 | ND |
| *Marseilleviridae* | Marseillievirus | H2A, H2B, H4, H3 | MC1* |
| *Mamonoviridae* | Medusavirus | H2A, H2B, H4, H3, H1 | ND |
| ***Mimiviridae*** | **APMV** | **ND** | **MC1** |
| *Phycodnaviridae* | Dishui Lake Phycodnavirus 1 | H3# | MC1, DVNP |
| *Poxviridae* | Vaccinia Virus | ND | ND |
| *Pandoraviridae* | Pandoravirus salinus | H2B | ND |

ND: not detected. * - metagenome assembled; # - a histone variant.
